# Pear (*Pyrus communis* L. cv. Conference) has shade-tolerant features allowing for consistent agrivoltaic crop yield

**DOI:** 10.1101/2024.04.24.590973

**Authors:** Thomas Reher, Brecht Willockx, Ann Schenk, Jolien Bisschop, Yasmin Huyghe, Bart M. Nicolaï, Johan A. Martens, Jan Diels, Jan Cappelle, Bram Van de Poel

**Affiliations:** Molecular Plant Hormone Physiology, Division of Crop Biotechnics, Department of Biosystems, KU Leuven, Willem de Croylaan 42, 3001, Leuven, Belgium; Electrical Energy Systems and Applications (ELECTA) Ghent, Department of Electrical Engineering, KU Leuven, Gebroeders De Smetstraat 1, Ghent, 9000, Belgium; VCBT - Flanders Centre of Postharvest Technology, Willem de Croylaan 42, 3001 Leuven, Belgium; MeBioS, Department of Biosystems, KU Leuven, Willem de Croylaan 42, 3001 Leuven, Belgium; Centre for Surface Chemistry and Catalysis: Characterization and Application Team (COK-KAT), Department of Microbial and Molecular Systems, KU Leuven, Celestijnenlaan 200F, box 2461, 3001 Leuven, Belgium; Division of Forest, Nature and Landscape, Department of Earth and Environmental Sciences, KU Leuven (University of Leuven), Celestijnenlaan 200E, 3001, Leuven, Belgium; KU Leuven Plant Institute (LPI), KU Leuven, Kasteelpark Arenberg 31, 3001 Leuven, Belgium

**Keywords:** agrivoltaics, agronomy, AV, light adaptation, shading, renewable energy, crop performance, fruit quality

## Abstract

Transitioning to a fossil fuel free society requires an increase in solar energy production. However, expanding solar power to farmland competes with food production. Additionally, climate change threatens food security and leads increasingly to yield losses.

Agrivoltaics (AV) systems produce solar energy and food on the same field, while sheltering crops. In AV systems, crops grow in a modified environment with reduced solar irradiance, a tempered microclimate and a potential physical cover protecting against hail damage.

This research describes pear production under an AV pilot with 24% light reduction for 3 consecutive seasons. AV pear trees yielded 14% less than the reference. Flowering and fruit set was unchanged while AV reduced leaf flavonoid levels. The leaf photosynthetic light response was identical, yet a delayed leaf senescence under AV suggests an adaptation to the modified environment. AV impacted fruit shape, as there was an increase in the number of bottle shaped pears and a reduction in caliber. Other fruit quality traits were broadly unaffected, yet postharvest ethylene production was higher for AV fruit in 2022 than for the control.

This study demonstrates that AV systems hold potential for pear production under temperate climates and highlights plant adaptations that make this possible.

**Highlight:** Pear cultivation in agrivoltaic systems integrates renewable energy and sustainable fruit production. This study provides insights into crop yield, fruit quality, and plant adaptation towards an agrivoltaic environment.

## Introduction

Pome fruit, and especially pear hold an important place in the high-value fruit market. While apple production remains more widespread worldwide; pears are the most important pome fruit in Belgium, with an annual production value of € 227 million (Departement Landbouw en Visserij, 2023). Changing climate and increasingly tight margins put pressure on farmers to innovate, making pear production increasingly reliant on technical support systems such as irrigation or protective coverings.

Agrivoltaic (AV) production systems combine solar energy generation and agriculture on the same land and can offer physical cover to crops while generating additional yield from energy production, ultimately raising farmer income. AV systems present opportunities in achieving the current, and future climate goals (Proctor *et al*., 2021) but at its inception, agrivoltaics (AV) served principally as a tool to make use of the neglected space between ground-mounted solar photovoltaic (PV) installations (Goetzberger and Zastrow, 1982). Recently the focus of AV shifted to elevated PV modules augmenting agriculture, rather than crops filling out a solar power plant’s unused space (Dupraz *et al*., 2011). This shift towards a crop focused integration of renewable energy with agricultural production better addresses the current needs for both clean energy and sustainable food production. Unlike open-field agriculture, orchards typically do not employ crop rotation, rely on smaller agricultural vehicles, and often already implement some form of physical protection such as polythene film or hail net covers. Fruit orchards which, unlike annual crops, have a production lifespan of several decades, offer a unique context for AV. Well-designed AV systems can serve as a protective structure against adverse weather by modifying crop microclimate, reducing water evaporation from soil or mitigating extreme temperature impacts on crops (Barron-Gafford et al., 2019; Juillion et al., 2022), and can produce more consistent yields in a changing climate (Pataczek *et al*., 2023). On the other hand, AV systems always reduce irradiance, impacting yield and productivity of crops. Despite this, AV has the potential to achieve an increased land-use efficiency by combining energy and food production.

Early explorations of crop responses to AV focused on leafy vegetables and field crops (Weselek *et al*., 2019). Other research focused on crops in areas with Mediterranean climates or grapevine production in areas where the higher levels of irradiance may cause undesirable fruit sugar levels (Padilla *et al*., 2022). However, it remains largely unclear how AV systems can benefit perennial crop systems, particularly in temperate regions susceptible to variable weather conditions, with limited irradiance and other climatic challenges. In Germany, new experimental AV sites have been constructed with apples (Kressbron, Bavendorf and Gelsdorf), where multiple cultivars will be examined (Trommsdorff *et al*., 2023). In Llupia (France), a new plantation of pear will explore AV under a Mediterranean climate. The actual impact of AV on mature pear fruit production remains unknown so far.

Previous research studied pear productivity under homogeneous shade of hail-netting structures, which serve as a protective measure against physical damage. While they are an effective tool in managing hair damage to fruit (Kiprijanovski *et al*., 2016), these nets represent a significant cost to the grower and produce between 8-15 % shade. Under uniform shade netting, Peavey et al. (2022) showed a mixed response of pears at 30 % shading in Australia, with modified appearance but similar firmness and overall yield. Garriz et al. (1997) reported a decrease in pear fruit number but increase in firmness under shade netting in Argentina. Kappel and Neilsen (1994) related fruit number and quality to the % of sky coverage. Miller et al. (2015), showed that, while total time of shade had the greatest influence on yield, shading of apple in the morning rather than afternoon, had negative effects on photosynthetic assimilates. The spatio-temporal shading pattern produced by elevated AV systems, however, differs greatly from the more homogeneous light reduction from netting.

The agrivoltaic environment is characterized by periodic blocks of shade occurring in an intermitted timeframe, depending on system design and orientation (Valle *et al*., 2017). This pattern of shading, where sharp increases and decreases of light intensity alternate, results in more variability than that of shade nettings but more closely approaches the natural shading dynamics in a canopy. Shading due to transient cloud cover or from other leaves occurs in a much more erratic way. Plant photosynthesis responds dynamically to adapt to these variable light conditions (Kaiser *et al*., 2015, 2018). Within a tree canopy, sunflecks are a critical aspect of the dynamic light environment. They are characterized by brief, intense periods of sunlight that penetrate through gaps in a canopy or cloud cover, illuminating the understory or ground. Sunflecks can vary in duration from a few seconds to several minutes, and their frequency, duration, and intensity can significantly influence the photosynthetic efficiency of plants adapted to such fluctuating light environments (Salter *et al*., 2019). Long et al. (2022) underscored the importance of understanding plant responses to intermittent shading caused by sunflecks. If a leaf experiences high light exposure before brief shading, it responds better to subsequent sunflecks. In essence, induction during one sunfleck primes the leaf for better utilization of subsequent sunflecks (Way and Pearcy, 2012). As such, an AV system might mimic the dynamic light environment of sunflecks experienced in a tree canopy, but in a different time dimension.

For crops grown under AV systems, understanding the optimal balance of light and shade can help maximize photosynthesis and yield. Also, additional benefits of the protective cover and altered microclimate could secure yield and fruit quality. In this work we examined how pear trees (*Pyrus communis* L. cv. “Conference”) respond to fluctuating shade across three consecutive growing seasons in an agrivoltaic pilot installation under a temperate maritime climate in Belgium. We observed a consistent decrease in fruit yield with 16 %, but little differences in postharvest fruit quality when compared to a hail-netting control. The crop light environment was found to differ from the control both in intensity as well as in spectral composition under the semitransparent modules. Additionally, the partially covered system resulted in a modified microclimate, but not impacting flowering phenology and fruit drop progression. Despite exhibiting a comparable light response of leaf photosynthesis for both AV and control trees, we recorded a number of physiological adaptations to shade as a result of the agrivoltaic environment.

## Materials and methods

### Field plot layout

The pear orchard is located on a commercial farm in Bierbeek, Belgium (50°49’05.6“N 4°46’34.5”E) and equipped with an AV system as presented in Willockx et al. (2024) (Fig. 1). Bisol Lumina 185 Wp (1×1.67 m) semi-transparent glass-foil monofacial PV modules were mounted on existing hail net supports in a double- landscape position at 4.2 m above ground-level. The modules feature a transparent backsheet (10 % light attenuation) and standard c- Si wafers (15.6 x 15.6 cm). Three rows of 21 m were placed immediately above the trees, of which only the middle row of trees was used for experiments (Fig. 1A), flanked by two buffer rows (due to boundary conditions). The field layout of the agrivoltaics and control plots are shown in Fig. 1B. To compensate for the lack of independent replicate plots for the AV treatment (due to construction limitations), the number of control plots was increased to five. In doing so, we can more accurately estimate variability present in the field and between trees. There is one agrivoltaics plot and five control plots distributed across the field, maintaining a buffer distance from the field edges and the buildings to the South. Each experimental plot is made up of 1 row of 12 trees. Control plots were covered by a crystal hail net throughout the season (∼9 % PAR reduction), which was removed from the AV plot.

**Fig. 1.**
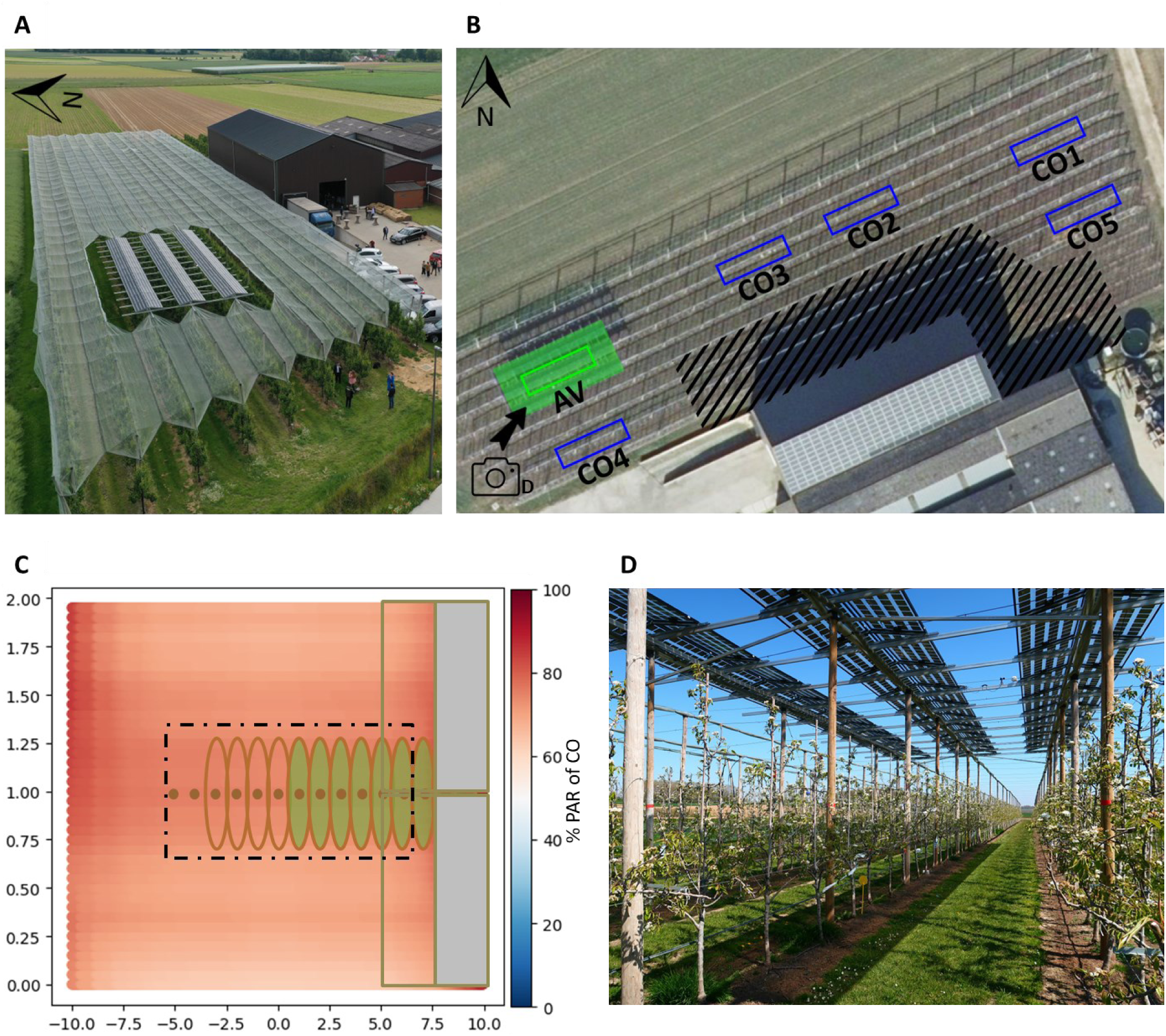
Agrivoltaic pear field lay-out. (A) Areal view of the pear orchard showing the agrivoltaics (AV) PV modules and the hail netting structure. (B) Aerial view of the plot layout for one AV (agrivoltaics) plot and 5 control (CO1-5; hail netting) plots. Each rectangle represents an experimental plot containing 12 pear trees. Area covered by PV modules highlighted in green, to illustrate the buffer area. Region impacted by shading from warehouse is delineated by shaded lines. (C) Schematic representation of the AV plot layout with simulated irradiance levels of photosynthetic active radiation (PAR) as compared to the control plots (hail net covered) according to Willockx et al. (2024)). Central 12 trees colored green, PV modules partially outlined in gray. Both trees and PV modules are continuous along the x axis but some are omitted here for visual clarity. (D) Picture taken in 2021 from the southern corner of the AV plot (indicated by camera icon on panel B).

This AV system resulted in a coverage ratio of 60 %, which combined with the transparency degree, transmits between 65 % and 72 % on average, of the incident photosynthetically active radiation on the ground plane depending on the time of year. The yearly average light reduction by the PV panels was 24 % photosynthetically active radiation (PAR) (Fig. 1C). Fig. 1D illustrates the shade at canopy level from semitransparent modules during early flowering.

### Orchard microclimate monitoring

Environmental data was collected with stand-alone temperature and relative humidity dataloggers (Testo 174H) mounted in custom made, 3d-printed weather shields and positioned in the middle and the base of the tree canopy. Other weather conditions were recorded using an on-site weather station (WH2600, Renkforce) and are summarized on a weekly basis in Supplementary **Error! Reference source not found.**.

### Light and spectral measurements

Incident PAR light was recorded using SQ500 sensors (Apogee) coupled to LogBox BLE multi-channel loggers (Novus Automation) for the base, middle and top of the tree canopy. The recorded values were averaged across the positions per timepoint in order to assess average PAR across the whole canopy. Light spectral composition was measured using a BLACK-CXR-SR-50 (StellarNet Inc.) spectroradiometer in the middle of the grass strip at 15 cm from ground level and in the tree canopy at 125 cm from ground level. Relative photon counts were normalized to the area under the curve of the PAR spectrum (400-700 nm). The transmission spectrum of the PV module backsheet was determined using the same equipment.

### Leaf temperature measurements

Leaf temperature was recorded using LAT-2B (Ecomatik) surface temperature sensor pairs positioned on and alongside the adaxial leaf surface, coupled to DL18 loggers (HOBO UX120-006M). Measurements were performed in duplicate in the center of the canopy for both AV and control plots. Recordings were made from 27/05/2022 until leaf drop at an interval of 1 minute.

### Phenology measurements

In January of 2021 and 2022, two random branches of each tree of each experimental plot were labeled and phenology was followed-up from the start of flowering. The predominant flowering stage for each flower cluster was recorded. Periodically throughout the flowering period, these flower clusters were assessed for development based on the classification presented by Claessen *et al*., (2022; see Supplementary **Error! Reference source not found.**). After flowering, the percentage of fruit drop was assessed weekly by counting the number of fruit remaining per cluster. Eventually a commercial manual thinning was performed by the grower leaving one to two fruit per cluster depending on tree load. In 2022, fruit clusters from marked branches were assessed for the presence or absence of apical flowers on June 10.

### Leaf pigment analysis

Leaf epidermis chlorophyll (Chl) and flavonoid (Flav) content were measured using a Dx4 (Dualex) leaf clip sensor (Cerovic *et al*., 2012). For this, 8 leaves per tree were selected at random throughout the canopy and measured. Measurements were performed regularly from 7/05/2021 until 15/11/2021 and from 28/04/2022 until 6/10/2022.

To determine total flavonoid content, the method of Cvek et al. (2007) was optimized for pear leaf samples. Leaves were snap frozen in liquid nitrogen and lyophilized (Beun – De Ronde, Bovenkamp, The Netherlands). After crushing, 125 mg frozen sample was mixed with 5 mL of 99.8 % ethanol. Thereafter the mixture was sonicated twice for 15 seconds at 100 J (Ultrasonic Liquid Processor, Fisher scientific). A standard curve of morin (≥ 95 %, Acros Organics B.V.B.A) was made using the same solvent. Thereafter 50 µL of morin solution was combined with 40 µL aluminum chloride reagent (2 % aluminum chloride and 5 % glacial acetic acid in methanol) in a cuvette and filled up to 1 mL with 5 % glacial acetic acid in methanol. Finally, the mixture was left for 30 min for the reaction to occur before the absorption was determined at 419 nm using a spectrophotometer (Shimadzu UV-1800). For leaf samples, 50 µL of the plant extract was combined in a cuvette together with 40 µL aluminum chloride reagent and filled up to 1 mL with 5 % glacial acetic acid in methanol and left to react for 30 minutes.

### Leaf biometry

Leaf area of at least 50 leaves per plot was determined using ImageJ (Schneider *et al*., 2012) from regular top-view images of individual leaves. Leaf thickness was measured from a small leaf section (central part) under an Olympus BX40 microscope equipped with a ToupTek Photonics E3ISPM camera module calibrated to a microscope grid slide (AV n=34, CO n=85).

### Leaf photosynthetic light response analysis

Photosynthetic light response curves were determined using an LCpro T advanced portable photosynthesis system (ADC Bioscientific LC6P) fitted with an external white light module at ambient [CO_2_]. Light response curves were made by measuring CO_2_ uptake rate at increasing light intensity up to 2175 µmol⋅m^-2^⋅s^-1^, waiting until equilibrium was reached between each increased step. The light response curves were fitted using a nonrectangular hyperbola- based model following Marshall and Biscoe (1980) using the “Photosynthesis” package (Stinziano *et al*., 2021) in R version 4.3.1 - “Beagle Scouts” and R Studio 2023.06.0 Build 421 “Mountain Hydrangea” (RStudio Team, 2020), and the dark respiration rate, the light compensation point and the carbon assimilation rate at saturating light levels were calculated. From the individual fitted curves, the light compensation point (I_max_) was determined using the tool developed by de Lobo et al. (2013) following Prioul and Chartier (1977).

### Fruit harvest and yield

The optimal harvest time was determined by the Flanders Centre of Postharvest Technology based on the time evolution of common maturity indexes (VCBT, Heverlee, Belgium) to be 13/09/2021, 24/08/2022, and 12/09/2023 respectively. Harvesting was done on two consecutive days each year. Per plot, pears per individual tree were harvested and the total number of pears per tree and total fruit weight per tree was quantified. A subsample of 12 fruit per tree was weighted to determine the average fruit weight.

### Fruit shape analysis

The diameter of the thickest part of the fruit was measured using slide calipers for 12 fruit per tree (n=144 per plot). Furthermore, pear shape for all fruit per tree was visually assessed at harvest and classified into one of two categories: normal or bottle-shaped as described by Smessaert et al. (2020). After harvest, commercial grading of the fruit was determined in bulk per plot using an optical sorting machine calibrated for Conference pears specifically (AWETA CSGM 3/14-R). Fruit were grouped in one of 8 categories: 15-45 mm, 45-50 mm, 50-55 mm, 55-60 mm, 60-65 mm, 65-70 mm, 70-75 mm, +75 mm. Total number of pears per category as well as an estimated weight per category was recorded automatically.

### Postharvest fruit quality and ethylene analysis

A subsample of 3 x 30 pears was taken from each plot. Of these, 30 fruit were analyzed at VCBT, immediately after harvest, for firmness using a benchtop penetrometer with a 0.5 cm^2^ plunger. Total dissolved solids were determined from the resulting juice using a digital refractometer (Atago, Tokyo, Japan), and reactive starch index determined after staining cut fruit with Lugol’s Iodine solution. The second batches of 30 fruit were placed in storage for 21 d at -0.6 °C and subsequently kept at 12 °C for 7 d to determine post-harvest shelf-life quality. The remaining 30 pears per plot were analyzed for their ethylene production rate. After storage at 3 °C for 14 d to induce climacteric ripening, individual fruit were weighed, labeled, and placed in a climate-controlled area at room temperature. Every other day, the fruit were placed individually in airtight containers (3.528 L) and left overnight to incubate at room temperature after which the headspace was sampled (1 mL). Samples were analyzed following Houben et al. (2022). Briefly, a 1 mL aliquot of headspace gas was injected into a Shimadzu gas chromatograph (GC-2014) equipped with a Porapak R50/80 stainless steel column (3 m x 3 mm) at an injector temperature of 150 °C. N_2_ was used as a carrier gas at 35 mL/min at a column temperature of 160 °C. Peaks were detected with a flame ionization detector at 250 °C. Ethylene concentrations measured in ppm and subsequently expressed as mmol g^-1^ h^-1^.

### Data analysis and statistics

Statistical analysis was performed using R version 4.3.1 - “Beagle Scouts” and R Studio 2023.06.0 Build 421 “Mountain Hydrangea” (RStudio Team, 2020). Data was visualized using the ggplot2 (Wickham, 2016), ggprism (Dawson, 2023) and export (Wenseleers and Vanderaa, 2020) packages.

To correctly assess variability in the trial design, a Linear Mixed-Effects Model was constructed using Lmer from the lme4 (Bates *et al*., 2015), with post-hoc testing using the lmerTest package (Kuznetsova *et al*., 2017) and Multcomp (Hothorn *et al*., 2008) packages. Ambient temperature differences between the treatments were assessed for normality using the Shapiro-Wilk test. If so, a one-sample t-test was performed to compare mean differences against a null hypothesis of zero difference (mu = 0). In cases of non-normality, the Wilcoxon signed-rank test was used instead.

All other parameters were compared by ANOVA or Kurskal-Wallis tests at p ≤ 0.05. Briefly, after an initial ANOVA, tests for normality of the residuals and for heteroscedasticity were done using the olsrr (Hebbali, 2020) and lmtest (Zeileis and Hothorn, 2002) packages. If the criteria for ANOVA were not met, a Kruskal-Wallis test was used. Pairwise comparison of means was done using Tukey HSD or Dunn post hoc tests (Kassambara, 2023).

## Results

### The AV system tempers the canopy microclimate

We observed that the agrivoltaic production system mildly changed the micro-climate of the orchard. Depending on the season and weather conditions, we noticed two temperature effects. First, crops under AV had a similar average daily temperature for most of the season when measured at the center of the canopy (Fig. 2A). However, within a single day, air temperature within the canopy differed more pronouncedly between AV and control plots. During early spring nights, when frost events occurred following moderately warm days (Fig. 2B), we recorded canopy night temperatures of approximately 0.5 °C warmer under AV compared to control plots, indicative of a slight frost protection (Fig 2C). Contrarily, during the daytime period, the average air temperature within the canopy under AV was slightly (0.36°C) lower compared to that of the control plots, but not leading to day-time frost events (Fig. 2C). Secondly, during hot summer days, the AV air temperature within the canopy seems to be similar to that of the control plots (Fig. 2D). A larger temperature variability is observed for differences in air temperature within the canopy in summer when comparing the AV site with the control plots (Fig. 2E). The temperature variability was less pronounced during the slightly warmer nights (0.5 °C) for AV plots.

**Fig. 2:**
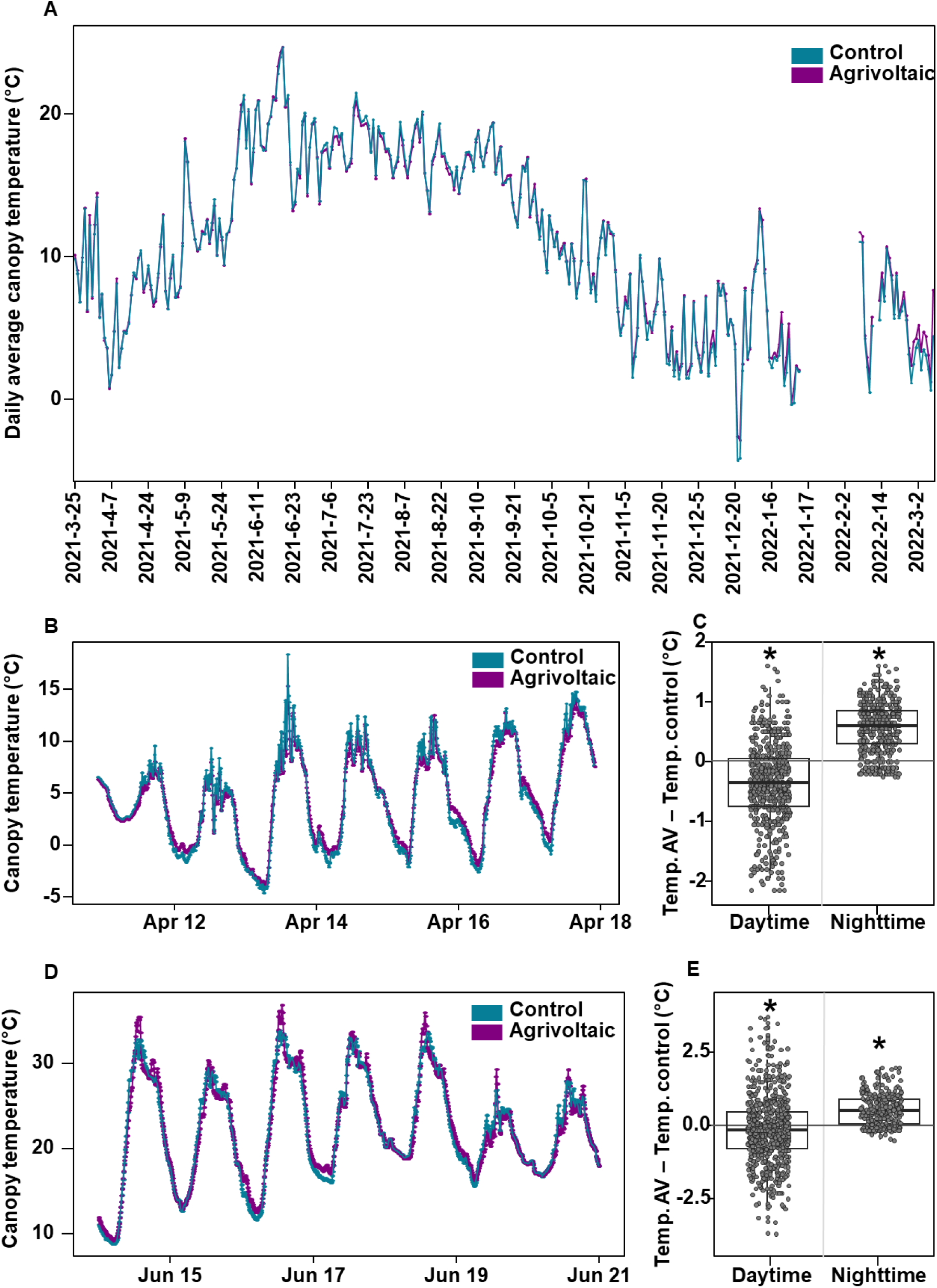
Seasonal effects of canopy air temperature for agrivoltaics and control plots. (A) The average daily temperatures for the 2021 season and its subsequent winter (2022). (B). Daily temperature profiles during several days of spring night frost. (C) Temperature AV minus temperature control for daytime (x̄=-0.364 °C) and nighttime (x̄=0.573 °C) during spring frost. (D) Daily temperature profiles during several days of a summer heatwave. (E) Temperature differential between control and agrivoltaics for daytime (x̄=-0.126 °C) and nighttime (x̄=0.540 °C) during the summer heatwave. Control (CO) in blue, AV in purple. Asterisks display significant differences (p < 0.05) between AV and control.

We wondered whether the subtle differences in canopy temperatures caused by the AV installation also impacted leaf temperature. Therefore, we measured the difference between the leaf temperature and the ambient temperature for the 2022 season and grouped them in ambient temperature blocks of 4 °C (Fig 3), both for nighttime and daytime periods. During the nighttime, leaf and ambient temperature are near identical for the control plots (an average differential equal to 0.05 °C; Fig. 3A). The agrivoltaic trees showed significantly slightly warmer leaves than the surrounding air during the night (an average differential equal to 0.15 °C), and this is consistent for the entire temperature range (Fig. 3A). Only for the hottest nights (24-27 °C), the control leaf temperature was on average 0.3 °C warmer (Fig. 3A). During the daytime, a more pronounced and significant deviation between control plots and agrivoltaic plots was visible. This deviation in leaf-air temperature difference between treatments increased when ambient temperatures were higher (Fig. 3B). Below 21 °C, daytime leaf temperature and ambient temperature under AV is nearly equal (an average differential equal to -0.02 °C), whereas the control trees showed a cooler leaf temperature than air temperature (an average differential equal to -0.46 °C). Above 21 °C, AV leaf temperatures increased (an average differential equal to 0.20 °C), whereas control leaves still remain cooler than their environment (an average differential equal to -0.46 °C). During hot summer days, the average agrivoltaics leaf temperature is almost 0.7 °C warmer than the surrounding air whereas control leaves remain close to their surrounding air temperature.

**Fig. 3:**
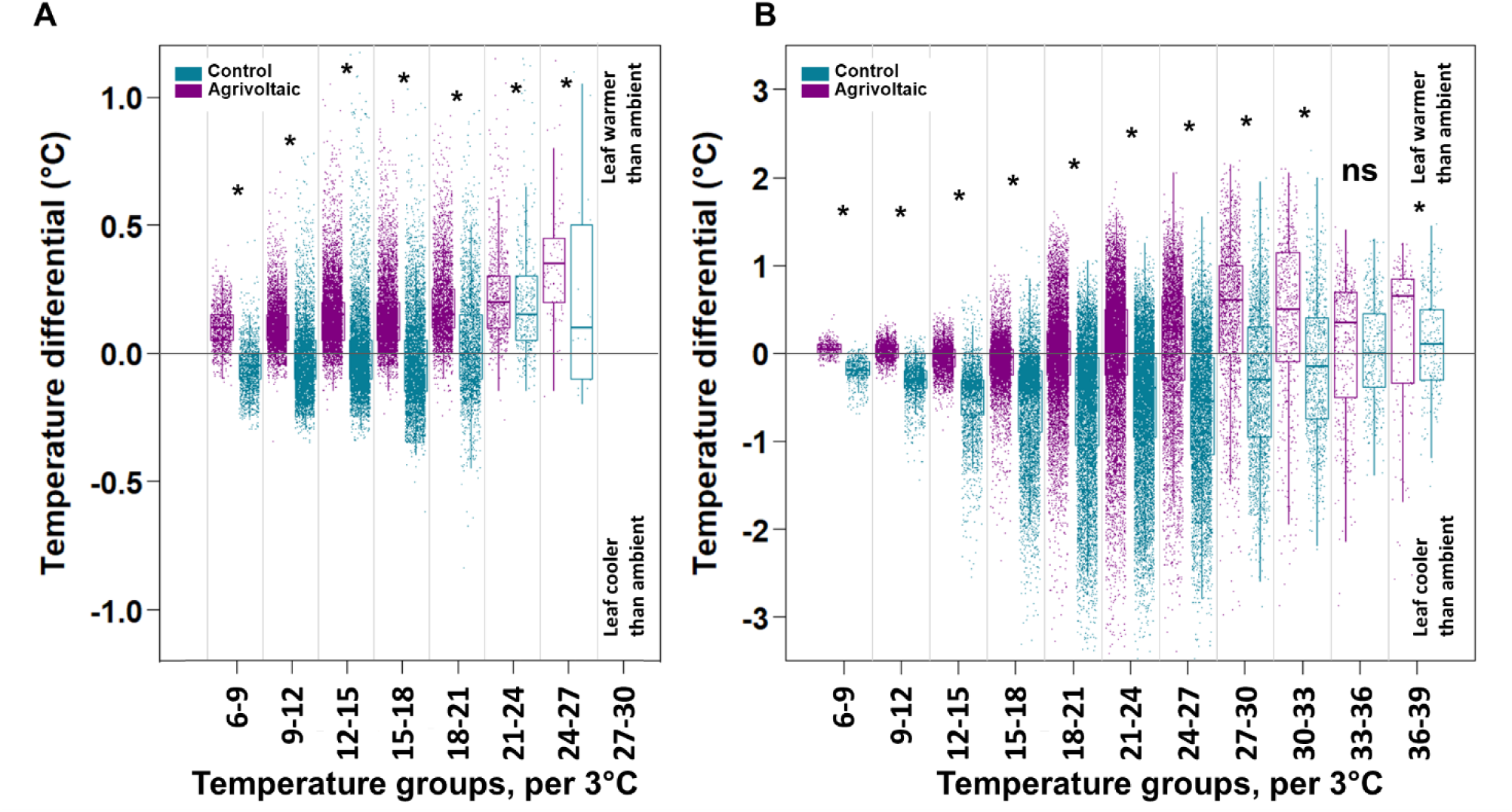
Pear leaf temperature dependency on different air temperatures. (A) Nighttime and (B) daytime leaf temperature minus ambient air temperature in its immediate vicinity clustered per temperature blocks of 3 °C for the 2022 season. Control (CO) in blue, AV in purple. Asterisks display significant differences (p < 0.05) between AV and control plots per temperature group. ns: non-significant.

### AV leads to light reduction and spectral changes

While light simulations offer valuable insights into the general light environment plants will register across the whole cultivation season (Fig. 1C), changes in canopy development are less easily accounted for *in silico*. Therefore, we measured the PAR, and observed an overall reduction of weekly PAR (daylight integral, DLI) under AV throughout the season (Fig.4A).

**Fig.4:**
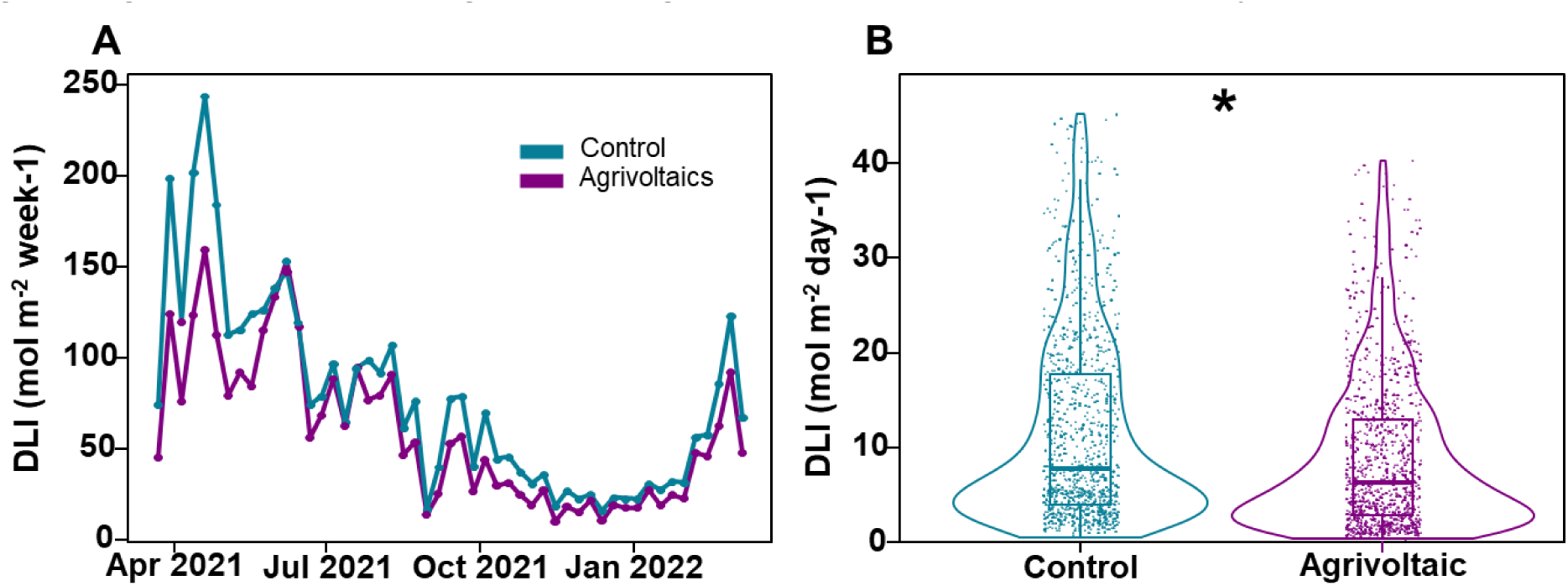
Agrivoltaics leads to a reduction in PAR light. (A) Profile of the weekly sum of daily light integral (DLI) at the pear canopy centerline. (B). Average daily PAR from three PAR sensors at top, center, and base of the canopy. Period: 26/03/2021-09/03/2022. Control (CO) in blue, AV in purple. Asterisks display significant differences (p < 0.05) between control and AV.

While the daily reduction percentage (23.8 %) remains reasonably consistent in periods of intense light, differences are more pronounced and can constitute up to 51 % DLI reduction. Correspondence of the measured PAR values to the light simulations of Fig. 1 are described in (Willockx *et al*., 2024). Fig.4b represents the average DLI values across the entire 2021 growing season, reflecting on average a 23.8 % reduction in canopy-perceived PAR.

Besides changing light intensity, the semitransparent glass-C-Si solar modules also influence the light spectral composition. Fig. 5 displays the changing light spectrum under AV when compared to the control area for two positions (in the tree canopy and in between tree rows), as well as the spectral transmissivity of the transparent PV module section (made up of glass, resin, and a diffuse plastic backsheet). Along the PAR spectrum, the PV modules absorb a relatively uniform fraction of light (around 10 %). However, in the UV-spectrum (300-400 nm) there is a 40 % reduction in UV radiation for the AV plots. We also observed a gradual decrease in irradiance in the blue spectrum (400-500 nm). No clear difference in spectrum was recorded between treatments for the longer wavelengths in the tree canopy. The red:far- red ratio remained similar between treatment at around 0.6. However, when measuring the red:far-red ratio for the grass-row in between the tree canopy, we observed a reduced R:FR ratio to about 0.3, again without differences between AV and the control.

**Fig. 5:**
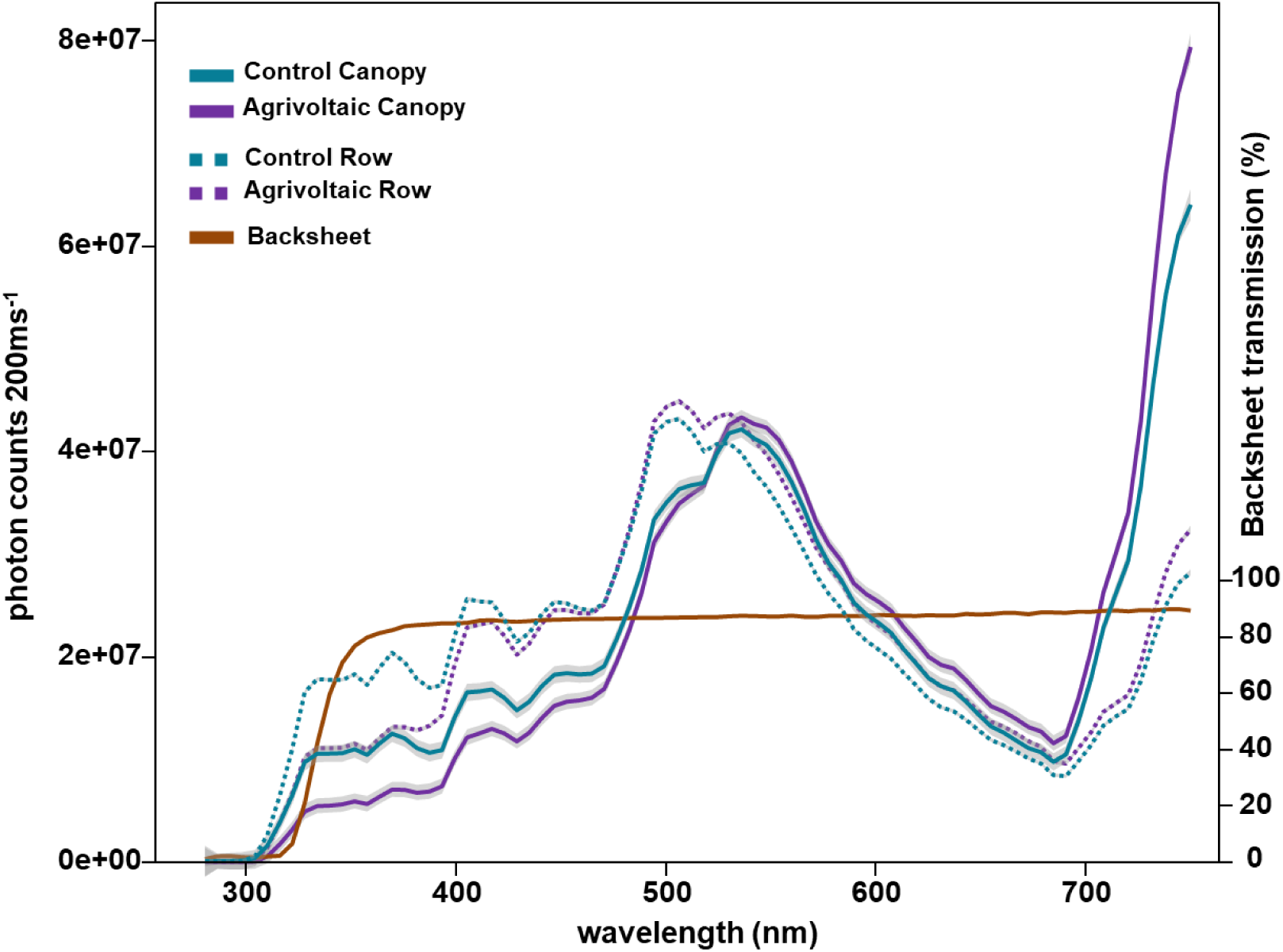
Normalized spectral composition of light under AV and in control plots and the PV backsheet transmissivity. Dotted lines recorded above the grass strip in between the tree rows. Solid lines recorded in the tree canopy. Blue: control plots (under hail-netting). Purple: agrivoltaic plot. Brown: relative light transmission of PV module backsheet. n=6. Spectroradiometer set at 200 ms integration time, 6 averaging passes per measurement.

### Photosynthetic light response remains similar under AV

Plants can alter their photosynthetic light perception efficiency when exposed to shade or altered spectral compositions (Liu and van Iersel, 2021). However, despite a clear drop in light intensity levels (on average 23.8 % PAR reduction) and spectral composition (Fig. 5), no differences in the photosynthetic light response curves were recorded between AV and control leaves. Fig. 6 visualizes the photosynthetic light response curves for pear leaves under AV and control conditions. It seems that AV leaves are slightly less efficient at 500 µmol m ^2^⋅s^-1^, but no significant differences are revealed for any of the photosynthetic parameters recorded: dark respiration rate (Fig. 6B), light compensation point (Fig. 6C), net CO_2_ assimilation rate at maximal photosynthetic irradiance (Fig. 6D), and light saturation point (I_max_, Fig. 6E). Despite not adapting to the AV system, pear tree photosynthesis appears to suffer equally little. This result is promising, given that in the fluctuating light environment, maximal utilization of direct light remains possible.

**Fig. 6:**
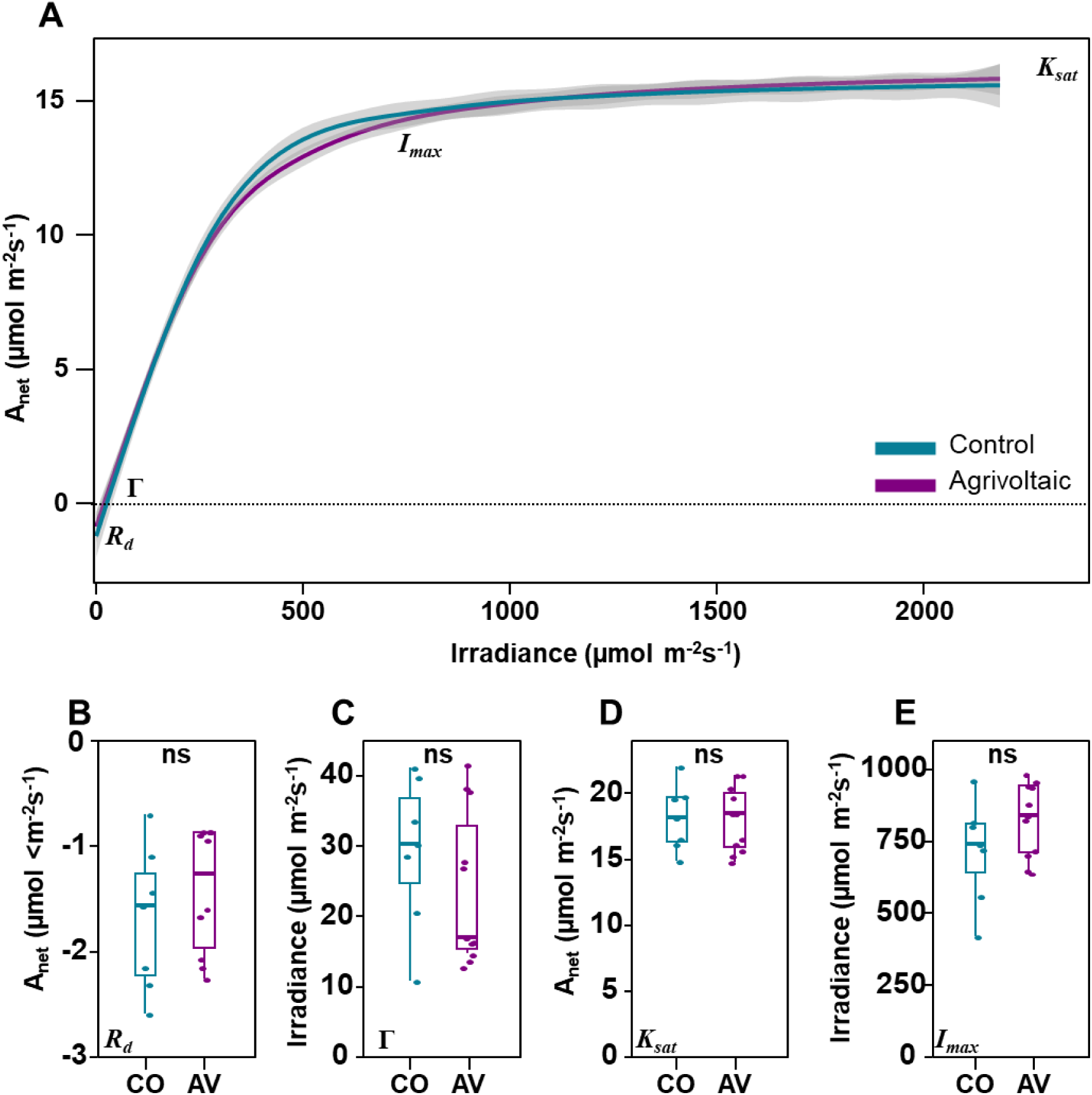
Photosynthetic light response curves for pear tree leaves under AV and for control plots. (A): Average of fitted light response curves with ± 95 % confidence interval. (B-E) Quantification of different photosynthetic performance indicators: (B) Dark respiration rate R_d_, (C) Light compensation point Γ, (D) Net CO2 assimilation rate at maximal photosynthetic irradiance K_sat_. and (E) Light saturation point I_max_. Control n=9, Agrivoltaic n=11. Control (CO) in blue, AV in purple. No significant differences (ns) were observed with p<0.05.

### AV induces subtle changes in leaf morphology

Beside photosynthetic adaptations, leaves exposed to shade can also adjust their leaf morphology (Wu *et al*., 2017). We observed that under AV, leaf thickness is slightly but significantly reduced (Fig. 7A). Individual leaf area was not significantly affected (Fig. 7B), but the specific leaf area under AV was recorded to be larger (Fig. 7C).

**Fig. 7:**
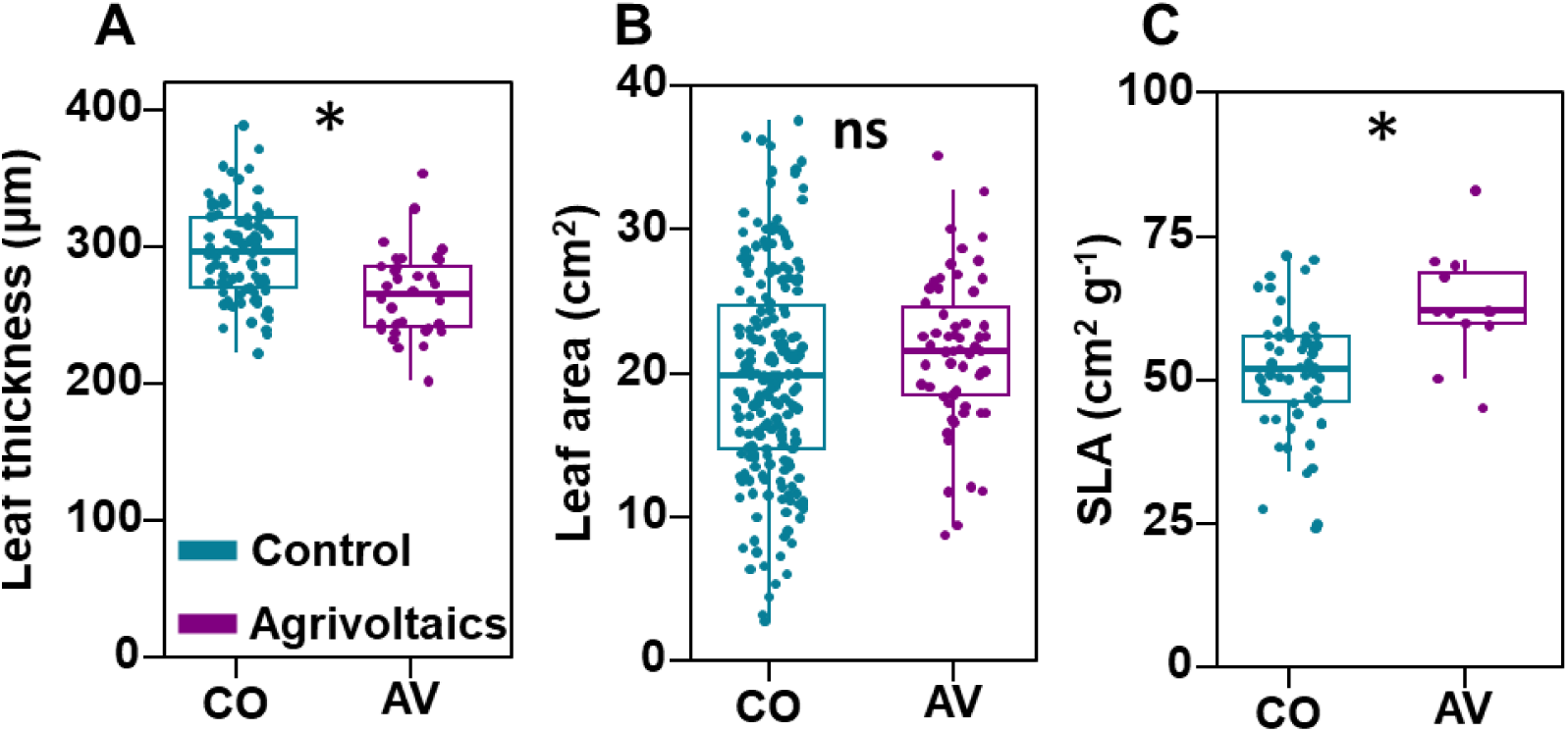
Agrivoltaics lead to subtle changes in leaf morphology. (A) Individual leaf thickness (CO n=85, AV n=34). (B) Individual leaf area (CO n=204, AV n=59). (C) Specific leaf area (SLA) (CO n=60, AV n=12). Data shown for the 2022 season. Control (CO) in blue, AV in purple. Asterisks display significant differences (p < 0.05) between AV and control. ns: non-significant.

### Leaf flavonoid and chlorophyll content is influenced by the AV conditions

An altered light intensity and spectrum, especially UV, can also affect leaf pigmentation (Brelsford *et al*., 2022). The leaf epidermis pigment content (Fig. 8) was measured with a non-destructive light transmittance device. Both chlorophyll (Chl) and flavonoid (Flav) values increased consistently throughout the season, peaking in late summer (Supplementary **Error! Reference source not found.**). Chlorophyll content was recorded to be slightly higher under AV in early season measurement both in 2021 and 2022 (Fig. 8A). However, this difference dissipated towards the late season. The leaf epidermis flavonoid measurements were significantly lower for AV compared to the control using the leaf clip method throughout the entire season for both years (Fig. 8B). When looking at whole-leaf flavonoid extracts, however, this difference could not be corroborated. The analytical measurements of total leaf flavonoid content showed no significant difference between AV and control leaves.

**Fig. 8:**
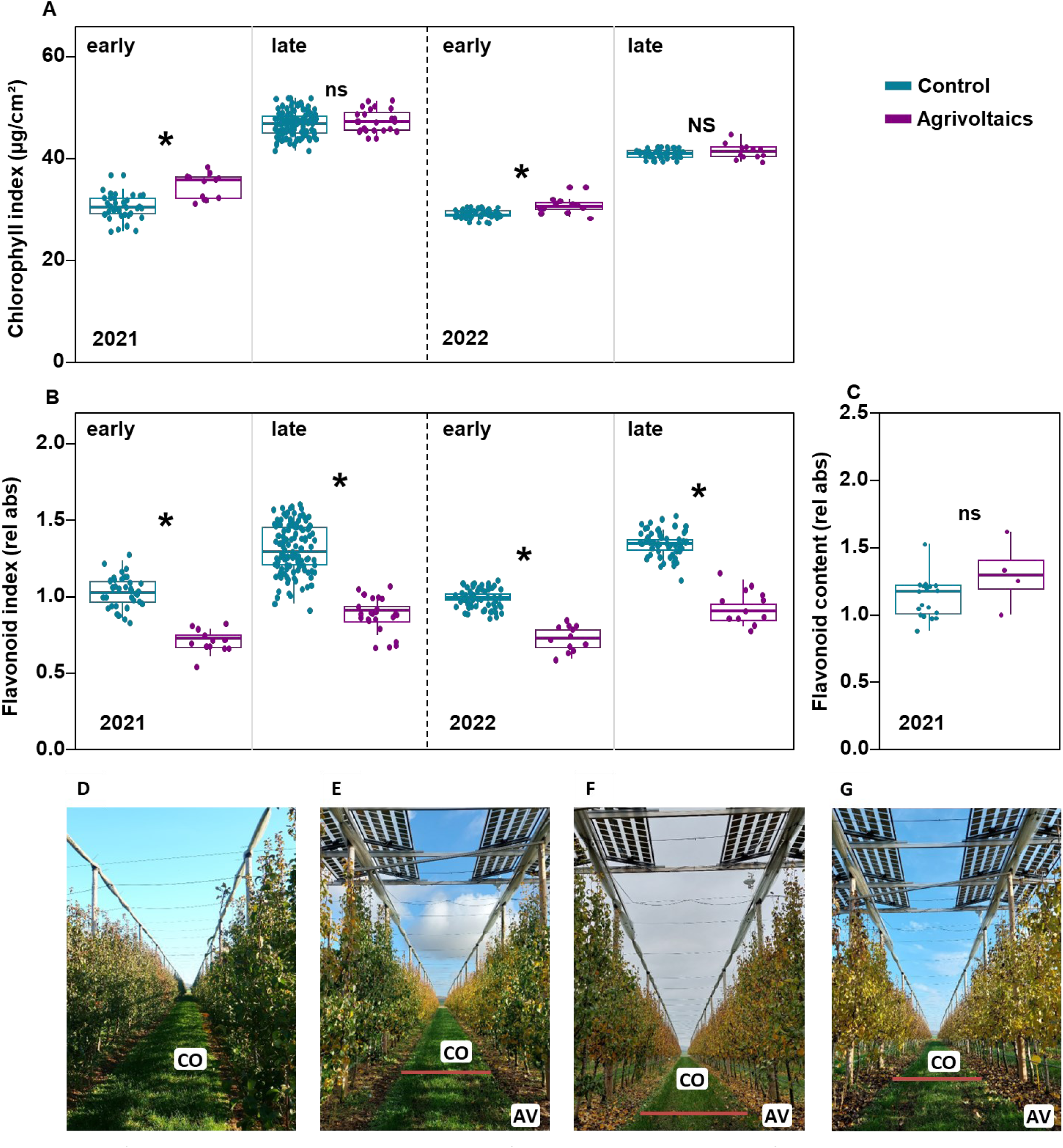
Pear leaf pigment changes under agrivoltaics. (A) Leaf epidermis chlorophyll content for 2021 early (10/5) and late (12/8) season and 2022 early (20/5) and late (11/8) season. (B) Leaf epidermis flavonoid content for 2021 early (10/5) and late (12/8) season and 2022 early (20/5) and late (11/8) season. (C) Total leaf flavonoid content for 2021 late season (12/8) analyzed using the AlCl2 method, optimized from (Cvek et al., 2007). 2021 early: CO n=36, AV n=12. 2021 late: CO n=120, AV n=24, 2022 early: CO n=60, AV n=12, 2022 late: CO n=56, AV n=12, Analytic: Control n=20, AV n=4. Control (CO) in blue, AV in purple. (D-G) Visual evolution of senescence in the row of the agrivoltaics (AV, forefront) and control (plot CO3, background) on (D): 27/10, (E): 5/11, (F): 11/11 (G): 18/11. No senescence visible for first date (panel D). Asterisks display significant differences (p < 0.05) between AV and control per timepoint. ns: non-significant.

Leaf clip measurements became unreliable in fall due to the occurrence of leaf senescence. A visual assessment of canopy discoloration during fall shows that AV trees retain their leaf chlorophyll longer, approximately 8-12 d depending on the year, compared to control trees Fig. 8 D-G which senescenced faster.

### Flowering and fruit development are unchanged under AV

In early spring, pear flower bud development occurs before leaves emerge. This period is key in determining yield potential, provided no unexpected damages occur. Spring frosts, as recorded in Fig. 2 can have devastating effects on flower viability and thus eventual yield. Despite small differences in the microclimate between the control and AV plants, we did not observe significant differences in flower development for both the 2021 (Fig. 9A) and 2022 season (Fig. 9B). Fig. 9 illustrate that the “balloon” stage of flower development progresses at a similar rate and time for both AV and control trees (for flower stage classification described in Supplementary **Error! Reference source not found.**).

**Fig. 9:**
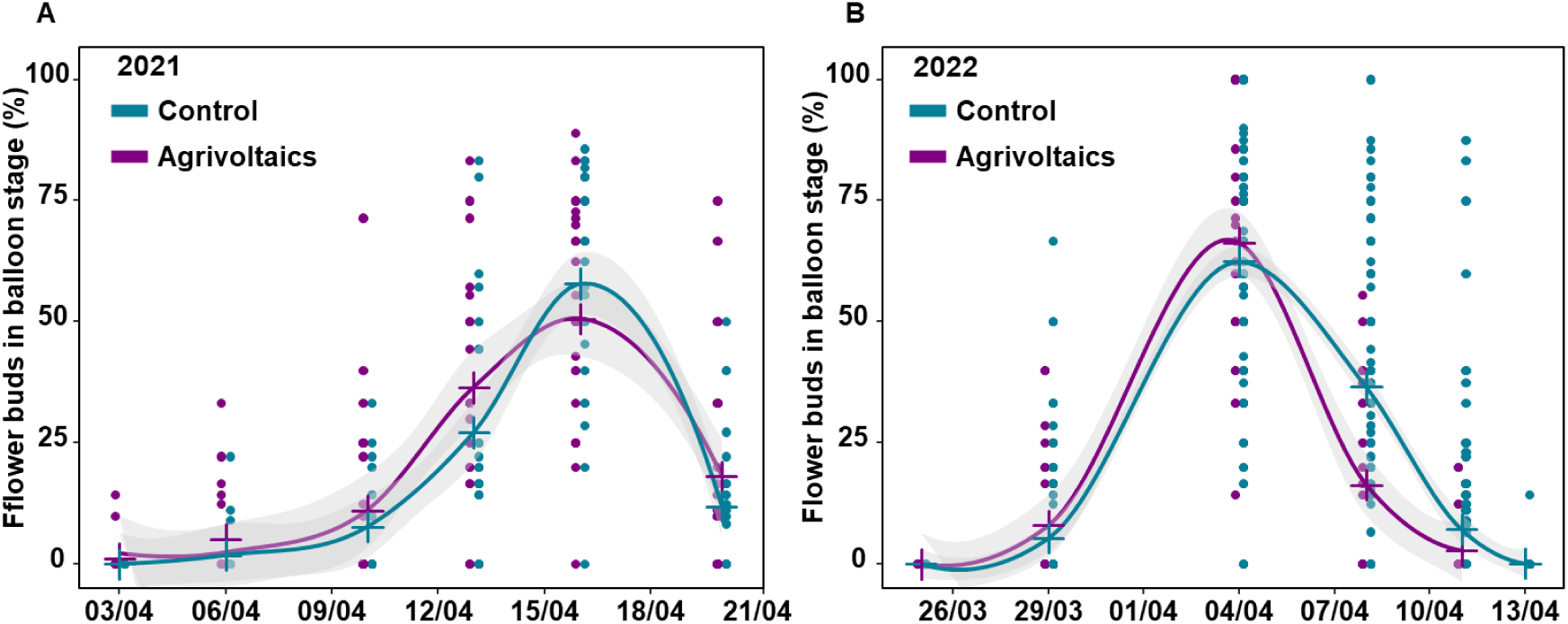
Progression of the occurrence of the balloon flower stage of pear trees. (A) 2021 season (B) 2022 season. Lines plotted using Locally Estimated Scatterplot Smoothing. n=variable with bud development. Control (CO) in blue, AV in purple.

Following pollination, a large number of the fruitlets that form initially will abort, leading to a reduced number of developing fruit per flower cluster. Fig. 10 shows the progression of fruit drop percentages during spring for the 2021 and 2022 season. Both the rate of fruit drop (Fig. 10A, C) and the final number of retained fruit (Fig. 10B, D) did not differ between AV and control trees, indicating that AV did not influence the fruit drop behavior of the tree. Note, that for both years, additional manual thinning was done after the end of the observations (as a commercial practice), reducing the number of fruit per cluster to 1.6 fruitlets on average.

**Fig. 10:**
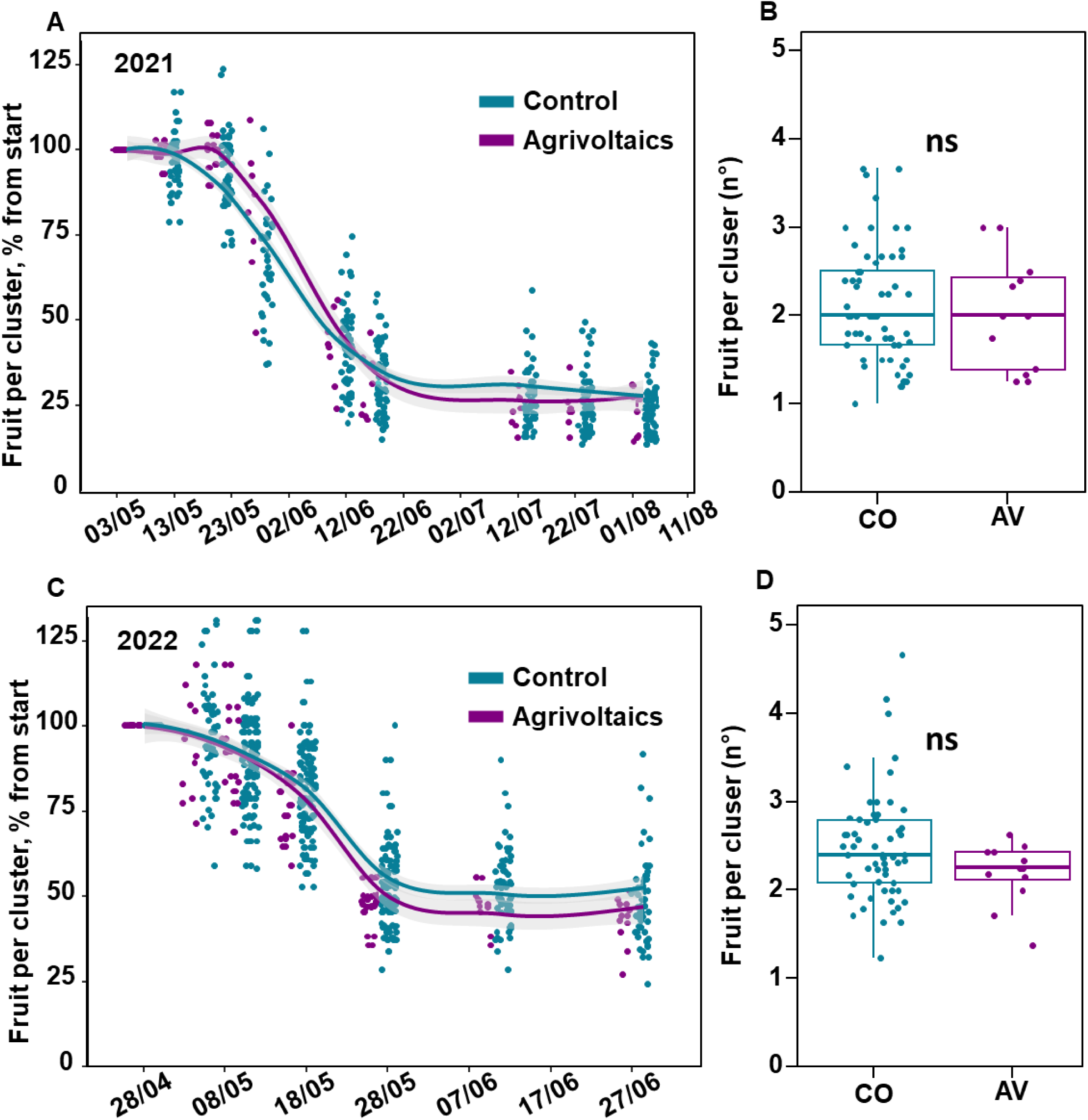
Evaluation of fruit drop progression under agrivoltaics. (A, C) Percentage of pear fruitlets remaining per flower cluster during the fruit drop period of the (A) 2021 and (C) 2022 season. (B, D) Residual number of fruits retained per flower cluster at the end of the fruit drop period for the (B) 2021 and (D) 2022 season. Control (CO) in blue, AV in purple. ns: non-significant for p<0.05.

### AV consistently negatively affects pear fruit yield

Across all three seasons, the average yield under AV was reduced by 15.0 % compared to the control plots (Fig. 11A). In 2021, yields dropped from 42.6 ton.ha^-1^ to 35.7 ton.ha^-1^ (16.2 %) under AV, in 2022 from 52.6 ton.ha^-1^ to 44.7 ton.ha^-1^ (15.0 %) and in 2023 from 47.6 ton.ha^-1^ to 41.1 ton.ha^-1^ (13.7%). Related to the individual fruit weight, only in 2021, a statistically significantly lower fruit weight (only 6.3 %) was observed under AV (Fig. 11B). For the other years, fruit weight nor fruit number per ha (Fig. 11C) were significantly lower for AV compared to the control. Fruit size, as measured by fruit diameter from manual subsamples, did not differ between AV and control for the three years (Fig. 11D). However, commercial grading of all fruit harvested per plot, demonstrated a slight shift in fruit caliber (Fig. 11E-G). There was a higher abundance of smaller fruit, and the fruit size distribution histogram shifted one category (5 mm) lower for AV plots compared to the control plots for 2021 (Fig. 11E) and 2023 (Fig. 11G), but not for 2022 (Fig. 11F).

**Fig. 11:**
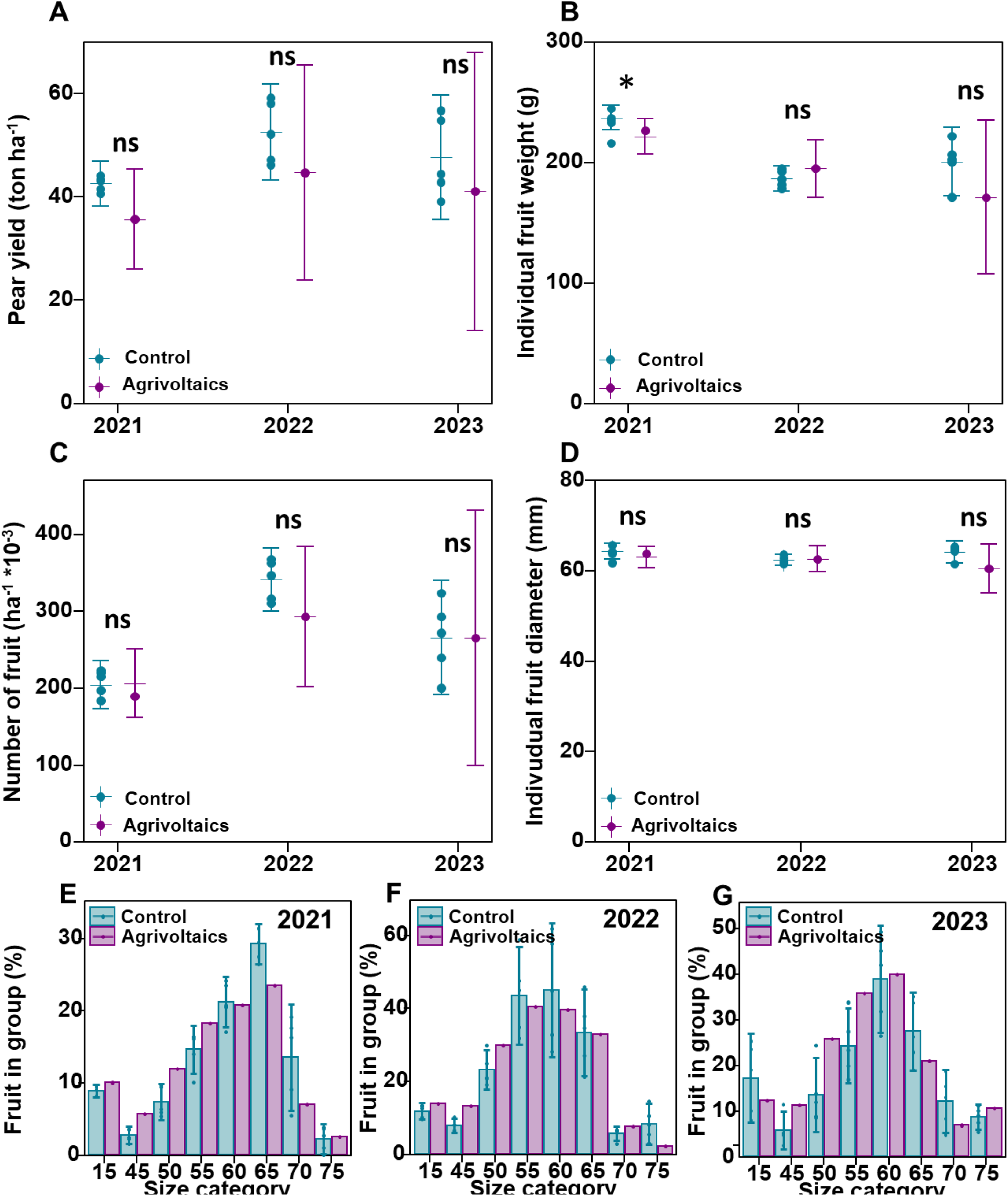
Impact of agrivoltaics on fruit yield and fruit characteristics. (A) pear fruit yield per hectare. Solid dots show field plot means, while the error bars show the treatment means with their 95 % confidence limits. (B) Average individual pear fruit weight. (C) Total pear fruit number per ha. (D). Average individual pear fruit diameter. Control (n = 5 plots), Agrivoltaic (n = 1 plot; based on all fruit harvested from 12 trees). (E-G) Commercial fruit size sorting per categories of 5 mm for the (E) 2021, (F) 2022, and (G) 2023 season. Group 15: undersized discards. Group 75: 75 mm and beyond. Control (n = 5), Agrivoltaic (n = 1; based on all fruit harvested from 12 trees). Control (CO) in blue, AV in purple. Solid dots show field plot means, while the error bars show the treatment means with their 95 % confidence limits Asterisks display significant differences (p < 0.05) between AV and control per year. ns: non-significant.

Fruit shape has an important influence on market value of the crop. We observed that the number of bottle-shaped pears was higher under AV compared to the control, which was significant for 2021. In 2022 and 2023 a similar trend was observed, but variability in the control plots was slightly larger (Fig. 12A). Bottled-shaped fruit are fruit that develop from the apical-flower of a flower cluster (Claessen *et al*., 2021). Therefore, we also quantified the fraction of flower clusters bearing apical flowers (for 2022) and found that agrivoltaic trees retain on average more apical flowers than control trees (Fig. 12B), which could explain the higher fraction of bottled-shaped fruit (Fig. 12A).

**Fig. 12:**
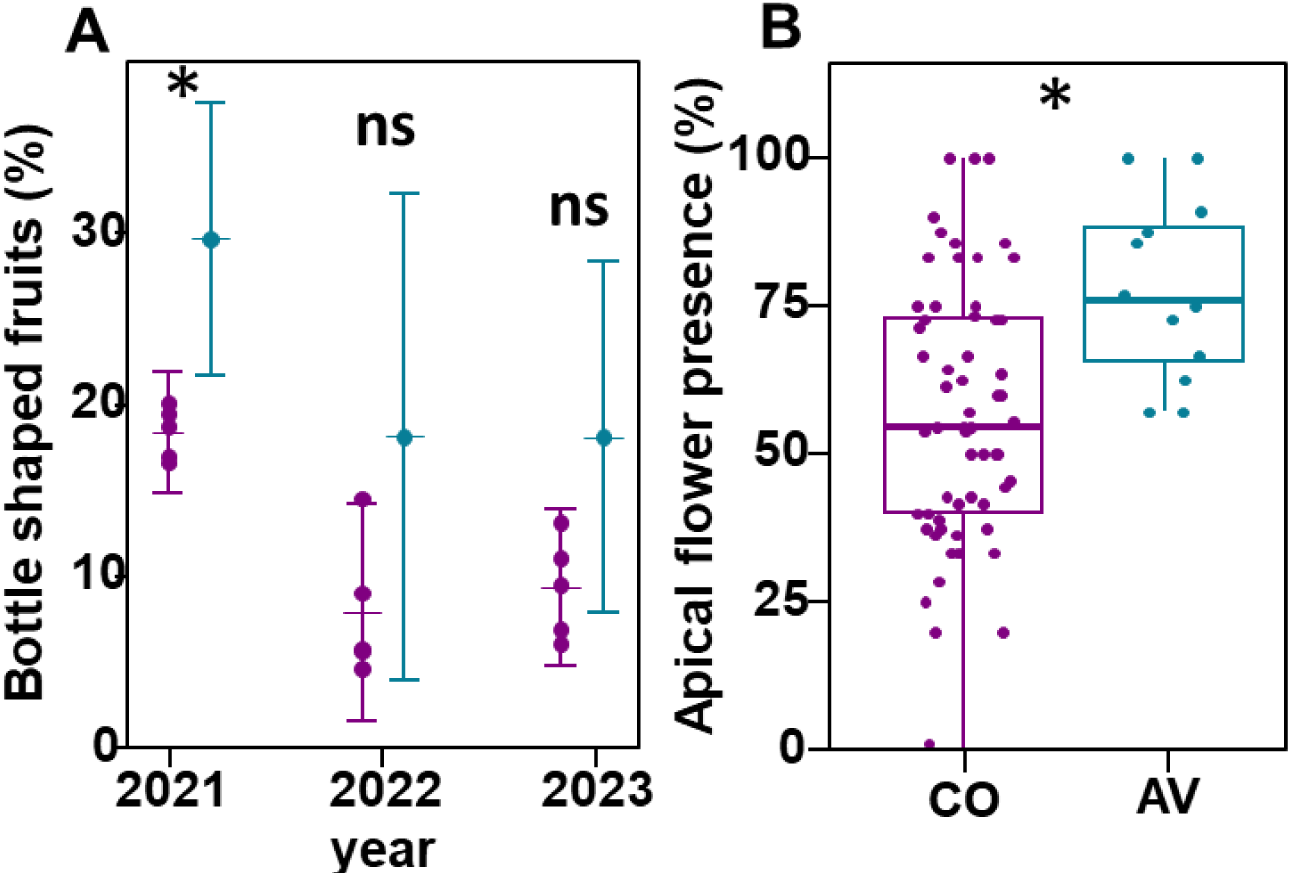
Percentage of bottle-shaped pears at harvest and presence of apical flowers in the field. (A) Percentage of bottle-shaped pears for 2021, 2022 and 2023 (Control n=5; AV n=1, based on all fruit harvested from 12 trees). Solid dots show field plot means, while the error bars show the treatment means with their 95 % confidence limits (B) Relative percentage of developing apical flowers retained per tree in 2022. Control (CO) in blue, AV in purple. Asterisks display significant differences (p < 0.05) between AV and control per year. ns: non-significant.

### AV does not influence fruit quality

After harvest, pears are typically chilled, and subsequently either brought to market to be sold or placed in modified atmosphere for long-term storage, depending on their physiological maturity (ripening stage). We assessed the ethylene production dynamics during post-harvest fruit ripening, to evaluate their climacteric behavior (Fig. 13) for two years (2021, Fig. 13A and 2022, Fig. 13B). We did not observe large differences in the ethylene production profile, suggesting that pear fruit harvested under AV ripen similar as control pear fruit. However, we did observe that AV pears reached a higher ethylene production level 3-4 days earlier during ripening for the 2022 season, indicative of a faster climacteric ripening process. We also assessed postharvest fruit quality parameters before and after 7 days of shelf-life storage (total dissolved solids (°brix); Fig. 13C&D; reactive starch index, Fig. 13E&F; and firmness, Fig. 13G&H). Only for 2021, post storage brix values were significantly lower for AV fruit. This did not affect the other quality parameters in 2021, neither before nor after 7 days of shelf life. Despite the higher ethylene production rate of 2022, none of the postharvest quality parameters were significantly affected.

**Fig. 13:**
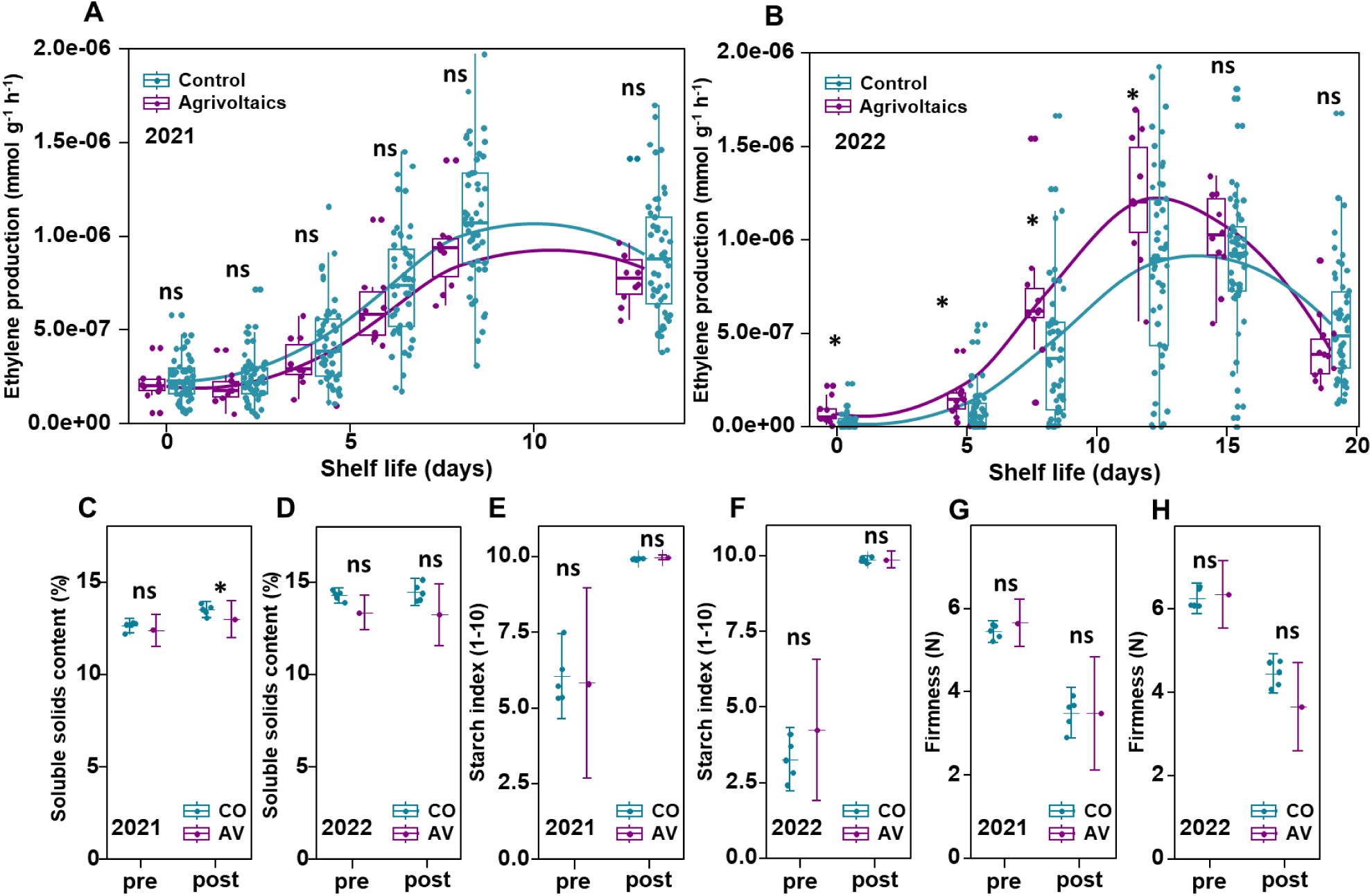
Postharvest quality parameters for the 2021 and 2022 harvests for control (CO) and agrivoltaic (AV) fruit. (A-B) Single pear ethylene production rates for (A) 2021 and (B) 2022. (C-D) Total dissolved solids, (E-F) reactive starch index, and (G-H) fruit firmness for the 2021 (C, E & G) and 2022 (D, F & H) season, before (pre) or after (post) a shelf-life period of 7 days at 12 °C. Control (CO) in blue, AV in purple. Asterisks display significant differences (p < 0.05) between AV and control per year. ns: non-significant. Panels A & B: agrivoltaics n=10 control n=50. Panels C, D, E, F, G & H: control n=5, AV n=1, based on all fruit harvested from 12 trees. Solid dots show field plot means, while the error bars show the treatment means with their 95 % confidence limits.

## Discussion

Pear trees, as well as many other fruit crops are sensitive to adverse weather such as spring frost, summer heatwaves and intense precipitation. Prolonged leaf wetness can also increase disease pressure and requires the application of additional agrochemicals. Global climate change causes an increase in extreme weather events, putting additional pressure on fragile agro-ecosystems. In order to safeguard crop productivity, growers make use of plastic covers or netting systems to mitigate some of these negative effects. This added level of control over the crop’s environment can help optimize productivity but comes at an additional cost. Agrivoltaic production systems can provide a similarly modified crop production system with the added benefit of sustainable energy production (821 MWh ha^-1^ y^-1^, (Willockx et al., 2024)). AV has the potential to serve as a competitive alternative for traditional crop protection systems and provides farmers with additional tools to increase climate resilience.

### The agrivoltaic microclimate reduces temperature extremes

Spring frosts are common during the flowering time of pear trees, typically occurring during the night when solar radiation is absent. The flowers, which develop before leaves emerge, are fully exposed to the elements, without protection. Growers can attempt to increase canopy temperatures during spring frost by lighting paraffin candles or using hot air blowers (Lakatos and Brotodjojo, 2022). Overhead irrigation during moments of freezing temperatures can also protect blooms from frost damage. Here, the latent heat released during freezing can keep the flower temperature above its damage threshold (Pan *et al*., 2024). Nevertheless, all these techniques are both energy and resource intensive.

In our field trials, frost damage did not occur, despite nighttime temperatures below zero during flowering. We believe this is caused by the increase in nighttime air temperature within the canopy under hail netting (control), which was more pronounced under AV, especially after sunny days (Fig. 2). The same effect was also observed by Lopez et al. (2023). They observed that this temperature difference at night was less pronounced when the weather was overcast, but also that nighttime temperatures remained higher overall due to cloud cover, posing less risk to frost damage.

Heat waves can be similarly detrimental to pear production. A tempering of ambient temperatures under AV was observed during summer heat waves. Our AV system mildly decreased extreme heat during the day. A similar decrease of peak temperature under AV on hot days has been shown in a Mediterranean climate for an apple orchard (Juillion *et al*., 2022). However, their system had a different mode of operation (dynamic tracking) permitting additional control over irradiance levels, whereas our design relied on a static PV system. While none of the fruit in the orchard, hail net control nor AV, suffered from sunburn, the neighboring fields without hail-netting protection were affected. We believe that the reduced irradiance of both the translucent low-density polyethylene (LDPE) fibers (crystal) netting (∼9 % reduce PAR levels compared to open air (Blanke, 2009)) and AV (24 % reduce PAR levels compared to hail nets) sufficiently reduced heat stress to mitigate sunburn damages. Netting has already been shown to be effective in offsetting sunburn (Iglesias and Alegre, 2006; Kalcsits *et al*., 2018). We now showed that static PV panels can have the same protective effect.

### Flowering and early fruit development is unaffected under AV

During three consecutive seasons, AV and control trees showed remarkably similar flower development. Despite the 2021 season’s start from buds initiated before the construction of the AV system, both in 2021 and 2022 the flower bud development was very similar (Fig. 9).

These observations contradict earlier findings for apple under AV (Juillion et al., 2022), which recorded a reduced flower bud development and a decreased fruit drop under AV, for a Mediterranean climate.

Whereas bud development determines fruit number at harvest, fruit size potential is largely influenced by cell division after pollination (Bound, 2021). According to Graham et al. (2021), pollinator behavior is unaffected by partial shade of PV systems. The similar levels of initial fruitlets after flowering for both AV and control trees leads us to believe that pollination was not an issue in our trials. Pear fruit drop has been shown to increase under shade (Einhorn & Arrington, 2018). However, we did not observe this effect under AV, recording similar fruit drop levels as for the hail-netting control. Hand thinning of excess fruitlets can be an effective tool in selecting for fruit size (Bound, 2021). Bearing in mind the relation we observed between bottle-shaped fruit and apical flower position(Claessen *et al*., 2021) clever hand thinning may increase harvest quality significantly.

### Leaf pigment changes reveal an adaptation to the AV light environment

The AV plot experienced a 60 % reduced UV incidence. Pigments, especially flavonoids and flavonols are influenced by the level of UV radiation, as perceived by the UVR8 receptor (Meyer *et al*., 2021) and the cryptochromes (Brelsford *et al*., 2022). Flavonoids are photoprotective pigments and they help manage reactive oxygen species in plant tissue or contribute to free-radical scavenging (Goulas *et al*., 2004). The reduced flavonoid level under AV suggest that pear trees under solar panels experience less light stress and require lower levels of photo protectants than their control counterparts.

Autumn senescence in deciduous trees is triggered by decreasing daylength and a drop in temperature (Smart, 1994). Pear dormancy is principally induced by low temperatures (<12 °C), regardless of photoperiodic conditions (Heide and Prestrud, 2005). The slightly higher temperature (Fig. 2) under AV could thus explain the delayed senescence of ∼12 days (Fig. 8F-H) observed for AV compared to the control. Alternatively, a reduced UV-radiation beyond 350 nm can also influence the onset of senescence (Brelsford *et al*., 2022). Similar delays (14.3 days) were recorded for *Acer platanoides* under reduced UV exposure.

Chlorophyll content is known to increase with increasing level of shade, particularly for woody species (Poorter *et al*., 2019), in order to improve light capture (Sage and McKown, 2006, Kappel and Flore, 1983). Our measurements indicated a very limited increase in chlorophyll content for the early season measurements only, a trend that dissipated later in the season (Fig. 8).

### AV alters pear leaf morphology but not photosynthesis

A shaded environment influences leaf shape and morphology (Kim *et al*., 2005). Leaf size changes have been observed in many shaded cropping systems. Under AV, an increase in lettuce leaf size was observed by Marrou et al. (2013). Apple trees develop thinner leaves underneath dark hail nets (Solomakhin and Blanke, 2010). A similar effect was recorded for pear trees under shade netting in Argentina (Garriz *et al*., 1997). Interestingly, we observed some thinner leaves and an increased SLA for AV pear trees (Fig. 7), compared to the hail- net control. An increase in SLA has been described as an adaptation mechanism to more shaded conditions for many species and is said to play a key role in maximizing carbon gain at plant level (Evans and Poorter, 2001).

Plants exposed to fluctuating light also exhibit thinner leaves while maintaining photosynthetic rates comparable to those of plants subjected to constant irradiation, for the same leaf area (Vialet-Chabrand *et al*., 2017). Contrary to earlier findings for apple under AV (Juillion *et al*., 2022), we observed no significant changes in photosynthetic light responses under AV (Fig. 6). It is clear however that a large variability in the rate of photosynthetic induction after shade exists between species (Salter *et al*., 2019). Vialet-Chabrand *et al*., (2017) did note an increased photosynthetic potential of *Arabidopsis thaliana* under randomly fluctuating light compared to continuous lighting at identical DLI, hinting towards adaptation strategies towards dynamic light environments.

Durand et al. (2021) highlights the importance of sunflecks for the upper strata of a tree’s foliage. These periods of relatively brief, yet significant increases in irradiance are abundant in a leafy canopy. *Arabidopsis thaliana* has been shown to perform better under fluctuating light than at continuous intensity (Vialet-Chabrand *et al*., 2017) and upper tree canopies seem to benefit from long (>5 min) sunflecks (Durand *et al*., 2022).

Despite changes in leaf morphology (Fig. 7) and pigment content (Fig. 8), it is uncertain if our observations under the fluctuating light of AV results in a comparably increased photosynthetic potential at canopy level. Despite single leaf photosynthesis light response curves showing no difference between AV and control, we suggest studying whole-canopy photosynthesis in future work.

### Pear fruit yield is less for AV, yet consistent over 3 seasons

At an average light reduction level of 23.8 %, our AV plot produced on average 15 % less fruit yield while maintaining similar individual fruit weights and number (Fig. 11). This observation contradicts past shading research suggesting that overall yield could largely be maintained under limited shade thanks to increased fruit numbers (Kappel, 1989; Peavey *et al*., 2022).

The slightly inferior fruit caliber at harvest for the AV pears could in theory be compensated for by a later harvest, as described by Gim *et al*. (2020) for Asian pears (*Pyrus pyrifolia*) under AV. However, the higher ethylene production rate of AV fruit observed in 2022 leads us to believe that a delayed harvest for AV pears is not desirable.

Despite the economically acceptable yield loss (Fig. 11), the increase in the proportion of bottle-shaped pears, may be worrisome. Fruit shape is an important commercial quality attribute and bottle-shaped pears have a lower market value. Gibberellic acid (GA) sprays to increase fruit set can increase bottle-shaped pear numbers (Smessaert *et al*., 2020), but GA applications were the same for AV and control plots. We therefore believe the increase in bottle shaped pears to be related to the AV conditions. Claessen *et al*. (2021) described how bottle-shaped pears principally develop from the apical flower in the flower cluster. They describe how, when sufficient sink potential goes to the top fruit, these fruits do not abscise, and more of the misshapen fruit will be retained. Our observations of the AV treatment showed a higher percentage of apical flowers being retained, suggesting a shift in sink potential of the flowers occurs under agrivoltaic conditions.

### Conclusions and future perspectives for agrivoltaic pear cultivation

The potential for agrivoltaics to improve crop resilience, yield, and quality in the face of climate change is apparent and our positive results on pear AV present a compelling case for further research and innovation in this field. In the light of practical applications, the importance of social acceptance of AV systems for multiple stakeholders (Torma and Aschemann-Witzel, 2023) should be considered. Also, with commercial agrivoltaic ventures, care should be taken not to overemphasize the reliance on PV to make a profitable system. Trommsdorff et al. (2023) highlight how apple production under AV can be profitable, while reaping the benefit from additional PV income.

This study evaluated the implications of an AV set-up on pear crop performance, yield, and quality, making AV a suitable crop protection system with relatively little impact on fruit yield and quality, yet production additional sustainable energy. Despite our indications regarding changes in the crop protection due to microenvironmental effects, the actual AV system (radiation) energy balance and linked to that the soil water balance remain to be studied. Also, the impact of AV on pest and disease susceptibility remains to be explored. Future research on AV systems for perennial crops best entail multi-year trials to refine AV-specific effects on crop performance. This way, AV systems can be a promising tool to combine profits from solar energy while sustainably increasing crop resilience.

## Acknowledgements

We would like to thank Jan and Patrick Van Der Velpen for their support and input in managing the orchard. Thanks to Boerenbond for the drone shots. Also, thanks go out to all colleagues and friends who helped with the harvest.

## Author contributions

TR, BVDP: Conceptualization; TR: Data Curation; TR, BW: Formal Analysis; JM, JD, JC, BVDP: Funding Acquisition; TR, BW, AS, JB, YH: Investigation; TR, BW, JD, BVDP: Methodology; JC, BVDP: Project Administration; JD, BN: Resources; BW: Software; JM, BN, JC, BP, BVDP: Supervision; TR: Visualization; TR: Writing – Original Draft; BW, AS, JB, YH, JM, JD, BN, JC, BVDP: Writing – Review & Editing.

## Data availability

Data is available upon request.

## Abbreviations

AV: Agrivoltaic(s) CO - Control
Chl: Chlorophyll
DLI: Daylight integral
Flav: Flavonoids
Γ (Gamma): Light compensation point
GA: Gibberellic acid
I_max_: Light saturation point
K_sat_: Net CO2 assimilation rate at maximal photosynthetic irradiance
PV: Photovoltaic
PAR: Photosynthetically active radiation
R_d_: Dark respiration rate
LCOE: Levelized cost of electricity
SLA: Specific leaf area
UV: Ultraviolet
UV-A: Ultraviolet A (315-400 nm)
UV-B: Ultraviolet B (280-315 nm)
VCBT: Flanders center of postharvest technology
x̄: sample mean

